# Nucleic acid turnover and lipid remodeling distinguish T cell activation and exhaustion states via label-free Raman spectroscopy

**DOI:** 10.64898/2026.05.25.727733

**Authors:** Marissa Morales, Srilakshmi Premachandran, Shruthi Ravichandran, Loza F. Tadesse

## Abstract

T cell exhaustion impairs immune control of chronic diseases including tuberculosis, HIV, malaria, and cancer. Its clinical implications are vast, predicting HIV-associated malignancy and treatment response and limiting the efficacy of cell therapies. Despite the advantages of monitoring and removing exhausted T cells, current detection methods require expensive antibody labeling, destructive workflows, or days-long functional assays. Here, we introduce Raman spectroscopy as a label-free assay for distinguishing T cell states directly from culture while preserving viability for downstream use. We achieve >97% accuracy in discriminating unstimulated, activated, and exhausted T cells across three donors and multiple hardware setups. We identify vibrational modes associated with nucleic acid turnover and lipid remodeling as key features that distinguish T cell activation and exhaustion. In heterogeneous populations, we quantify exhaustion percentage with R^2^= 1 and strong correlation to adenine (r= −0.91) and amide II protein (r= 0.94) vibrational modes. This work establishes vibrational fingerprinting as a direct measure of T cell exhaustion beyond surface marker expression towards scalable immune diagnostics, in-line monitoring, and selective immunopheresis.

## Main

Chronic illnesses including tuberculosis, HIV, malaria, and cancer claim over 12 million lives annually.^1–4^ T cell exhaustion is a hallmark of disease progression in these conditions, where persistent antigen stimulation results in reduced effector function,^5^ increased expression of inhibitory receptors,^5^ mitochondrial dysfunction,^6^ and epigenetic reprogramming.^7^ This impaired immunity has numerous clinical implications across disease contexts and therapy efforts. In HIV, T cell exhaustion precedes the loss of viral control^8^ and predicts HIV-associated malignancy up to 12 months before diagnosis.^9^ In cancer, exhaustion signatures predict responses to chemo and immunotherapy,^10,11^ informing treatment selection. Within chimeric antigen receptor (CAR) T cell manufacturing, exhausted cells extend manufacturing timelines^12^ and increase production costs, while *in vivo*, exhausted CAR+ cells demonstrate reduced potency and persistence.^13–15^ Notably, therapy non-responders harbor 2.5-fold more exhausted CAR+ cells than responders.^15^ Thus, rapid identification of exhausted T cells is critical to improve patient stratification and disease monitoring and inform personalized treatment. Furthermore, selective removal of exhausted T cells holds potential to accelerate cell therapy manufacturing and introduce new therapeutic interventions for chronic disease, akin to emerging senolytic therapies that rejuvenate immunity by removing senescent T cells.^16^

However, the existing paradigm for analyzing T cell exhaustion is inadequate for these applications. Flow cytometry, the current gold standard, requires hours-long multistep labeling with expensive fluorophore-conjugated antibodies that are sensitive to photobleaching and degradation.^17^ These antibodies target user-selected surface markers that prevent unbiased molecular profiling. Antibody binding can also alter cell physiology and function, risking the potency and safety of cells in downstream applications.^18^ Available alternatives suffer their own limitations: RNA sequencing and mass cytometry employ highly destructive^19,20^ processes, rendering cells unusable for immunopheresis and therapy manufacturing. Cytotoxicity and proliferation assays require days-long culture,^21^ too slow to inform manufacturing or treatment decisions. In addition, while size-based separation rapidly distinguishes resting T cells from activated T cells^22^ without labeling, it cannot differentiate activated and exhausted cells due to their similar size. A label-free, non-destructive method capable of distinguishing exhausted T cells from unstimulated and activated cells remains an unmet challenge.

Here, we introduce Raman spectral fingerprinting of whole T cells as a label-free assay for distinguishing exhausted cells from activated and unstimulated cells directly from liquid culture. Based on the inelastic scattering of photons, Raman spectroscopy offers molecular sensitivity at the single-cell and subcellular level without antibody-dye labels, destructive processing, or prolonged functional assays. Prior work demonstrates that Raman spectroscopy with machine learning-based spectral analysis can identify immune cell types,^23,24^ monitor stem cell differentiation,^25^ and distinguish naive versus activated T cells.^26–30^ With advances in Raman-activated cell sorting,^31–33^ it holds potential as an in-line technology for dynamic monitoring and immunopheresis. To determine the feasibility of our approach, we induced T cell exhaustion via repeated bead-based antigen stimulation and verified upregulation of inhibitory receptors PD-1, LAG-3, and TIM-3^34,35^ by flow cytometry. Analyzing >7000 single cell Raman spectra from three donors across custom and commercial Raman spectrometers, we achieved >97% average classification accuracy using class labels defined by the current gold standard, flow cytometry. Feature extraction identified vibrational modes associated with nucleic acid turnover (788 cm^-1^ and 819 cm^-1^) and lipid remodeling (1260 cm^-1^ and 1457 cm^-1^) as discriminative signatures of exhaustion. Prediction of exhaustion percentage across 100 mixed cell batches with R^2^=1 demonstrated model generalizability to heterogeneous populations relevant to clinical samples, even when exhaustion is present at low abundance. The percentage of exhausted cells in a given batch was found to scale linearly with adenine- (730-735 cm^-1^) and amide II protein-associated (1559-1562 cm^-1^) vibrational modes with Pearson correlation coefficients of −0.91 and 0.94, respectively. Together, these findings demonstrate the ability of our approach to capture the metabolic alterations accompanying T cell exhaustion and differentiate this phenotype from unstimulated and activated T cells. This technique enables label-free assessment of T cell state with applications in immune diagnostics and treatment selection. With single-cell flow channels, this technique allows for label-free cell sorting towards improved cell therapy manufacturing and opens the possibility for new chronic disease treatments to restore immune function via the selective removal of exhausted T cells.

## Results

### Label-free Raman spectroscopy identifies T cell exhaustion

As summarized in Fig. 1, we induced activation and exhaustion in pan T cells using Dynabeads Human T Activator CD3/CD28. After allowing isolated cells to rest for one day, we plated a cell sample in fresh media without recombinant human interleukin-2 (rIL-2) in a quartz-bottom petri dish and conducted single cell Raman spectroscopy measurements (Day 0). The parent cell culture was then stimulated with fresh beads every four days for twelve days, repeating Raman measurements using samples at each time point (Days 4, 8, and 12). In parallel, we performed flow cytometry measurements, staining for well categorized inhibitory receptors PD-1, LAG-3, and TIM-3 as markers for T cell exhaustion,^34,35^ which allowed us to assign class labels at the population level. The single cell spectra and class labels were then used to train and evaluate a 1-D convolutional neural network (CNN) to predict cell state as either unstimulated, activated, or exhausted based on the highest probability score.

**Fig. 1:**
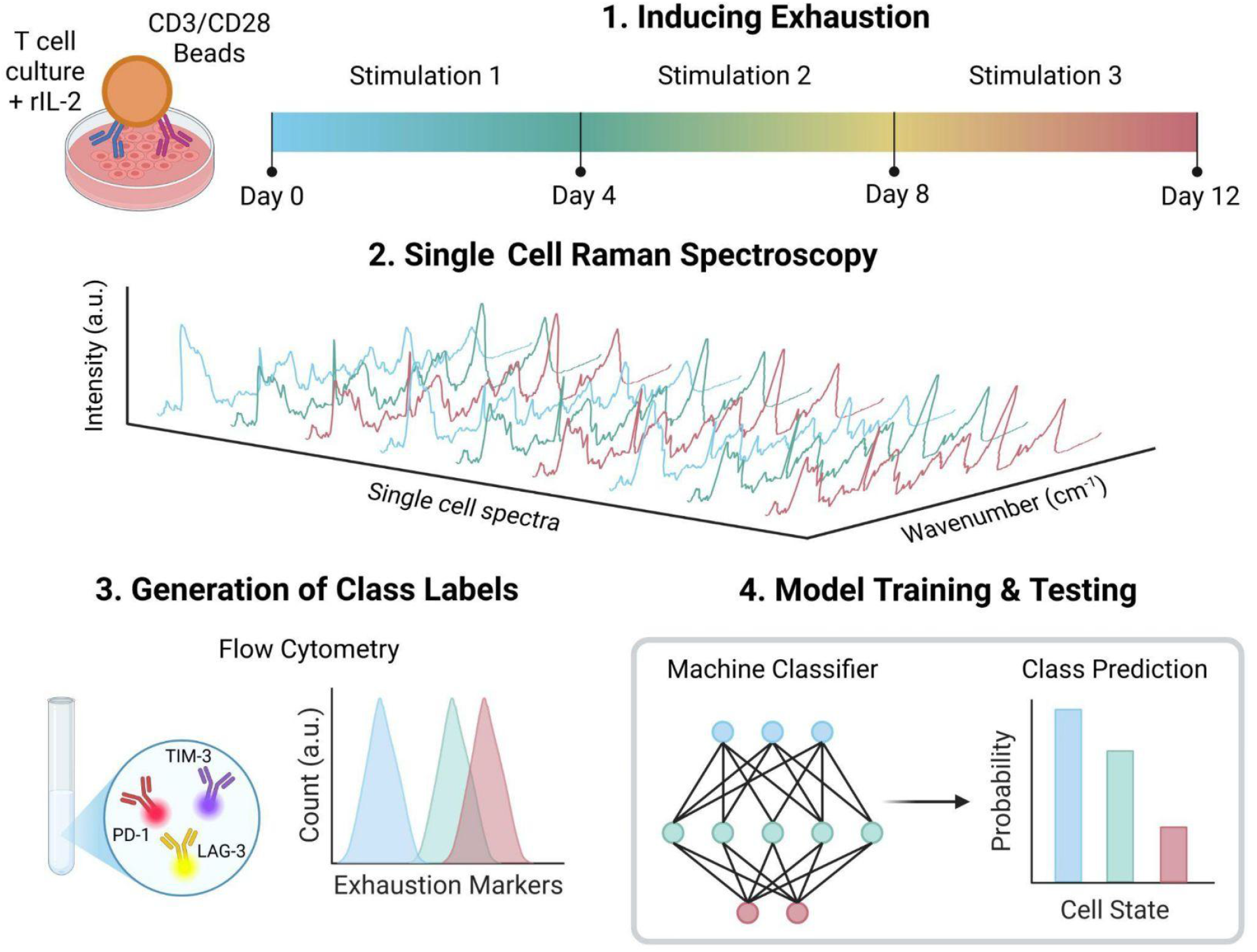
Schematic of label-free Raman identification of T cell exhaustion. To induce exhaustion, resting pan T cells were activated and re-stimulated every four days with Dynabeads Human T Activator CD3/CD28 for twelve days. At each time point, Dynabeads were removed (if present) and cells were plated into a quartz-bottom petri dish in fresh media without rIL-2. Single cell Raman spectra were measured, and signal intensities of exhaustion markers (PD-1, LAG-3, and TIM-3) were measured via flow cytometry to obtain class labels at the population-level according to cell state. The spectra were then pre-processed and input into a machine classifier with their corresponding class labels to train for and evaluate cell state prediction.

Flow cytometry analysis confirms that the T cells demonstrated an unstimulated phenotype at Day 0 with >94% of cells lacking all inhibitory receptors (Fig. 2A-B). Following the first stimulation at Day 4, T cells exhibited substantial upregulation of PD-1 and TIM-3 with moderate upregulation of LAG-3, consistent with prior *in vitro* models of T cell activation.^36,37^ After the second stimulation at Day 8, T cells further upregulated expression of LAG-3 and TIM-3 while maintaining expression of PD-1. After three stimulations at Day 12, upregulated expression levels of all three inhibitory receptors were maintained, with some CD8+ T cells gradually downregulating PD-1 (Fig. 2A) as previously observed in *in vitro* models of chronic stimulation.^37^ These results reflect the heterogeneity within exhaustion and further supports the need for more robust methods of staging exhausted T cells beyond surface marker expression. Taken together, we assigned class labels at the population level for Day 0, 4, and 12 as unstimulated, activated, and exhausted; Day 8 was identified as a transition state between activation and exhaustion and therefore left out of our three-state classification task. Gating strategy and flow cytometry results separated by donor are shown in Supplementary Figs. 1-4. Because CD8+ (Fig. 2A) and CD4+ (Fig. 2B) T cells exhibited similar inhibitory receptor upregulation trajectories and to better simulate conditions in real clinical samples, CD4+ and CD8+ T cells were not separated for Raman spectroscopy measurements. Independent Raman measurements with CD8+ T cells alone (Supplementary Fig. 5) confirm that classification is cell state dependent and not due to changes in CD4+/CD8+ ratios during stimulation and culture.

**Fig. 2:**
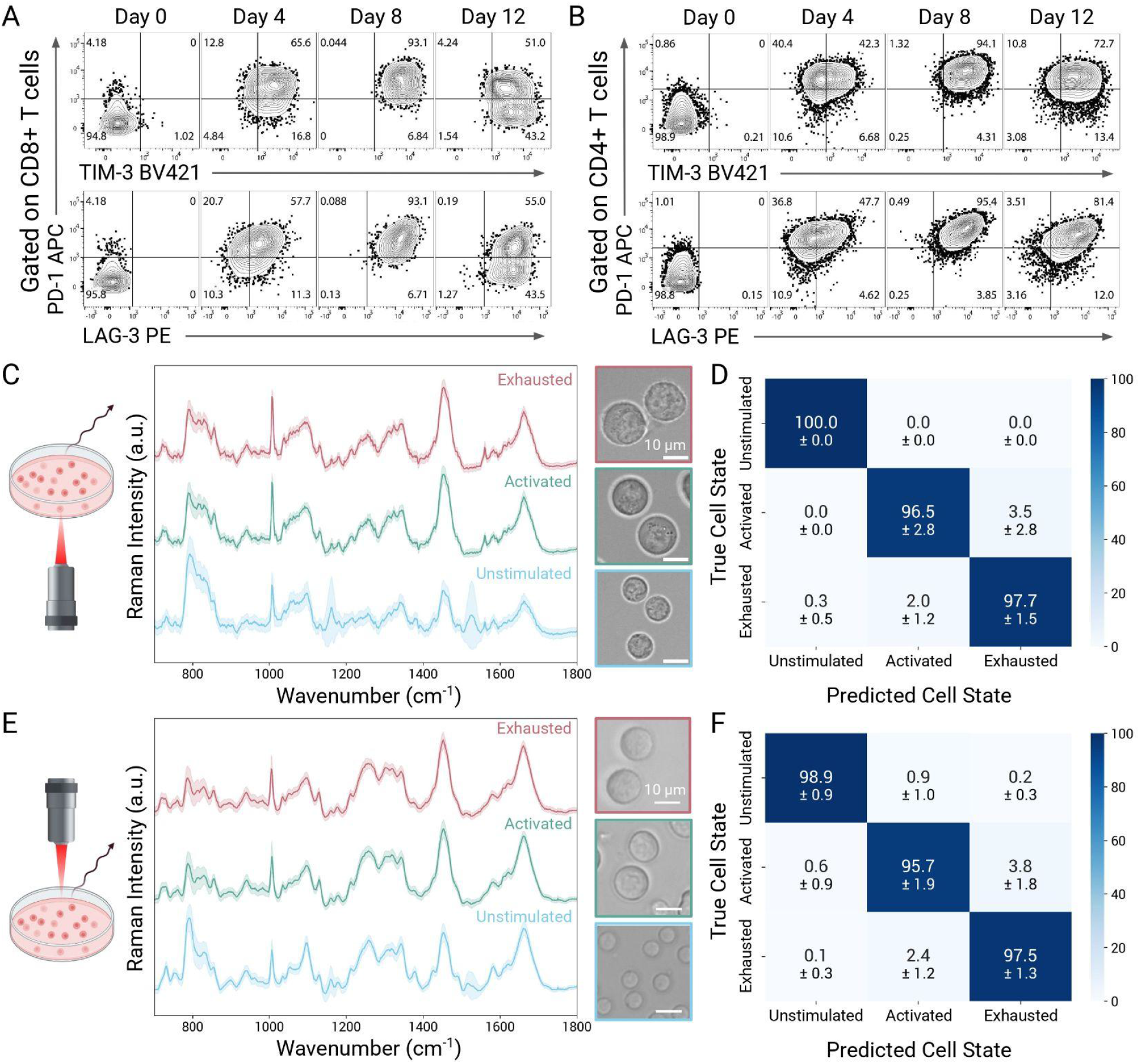
Raman spectroscopy distinguishes T cell activation and exhaustion. (A-B) Representative flow cytometry analysis of (A) CD8+ and (B) CD4+ T cells for the expression of inhibitory receptors PD-1, LAG-3, and TIM-3. (C-D) Identification via the inverted Raman system (1200 g/mm grating, multi-point scanning, spot size∼0.8 μm): (C) Average normalized single cell spectra from live T cells with representative brightfield images. Spectra are grouped by cell state with ±1 standard deviations (SD) shaded and used to generate the (D) normalized confusion matrix from our CNN. (E-F) Corresponding analyses to (C–D), conducted with data acquired from the upright Raman system (300 g/mm grating, single-point acquisition, spot size∼1.7 μm). Classifier performance in panels D and F was evaluated via stratified K-fold cross-validation using ten splits across three donors with three technical replicates each.

To determine whether T cell exhaustion could be identified without antibody labels, destructive processing, or functional assays, we collected >7000 single cell spectra (Fig. 2C & E) from primary T cells across three donors. Since instrument configurations vary across labs, we validated our approach using two independent Raman systems: a custom, inverted microscope (1200 g/mm grating, multi-point scanning, spot size∼0.8 μm) and a commercial, upright microscope (300 g/mm grating, single-point acquisition, spot size∼1.7 μm), both with 785 nm excitation. As expected, greater differences in cell spectra in the biological fingerprint region (700-1800 cm^-1^) are observed between unstimulated and activated cells than between activated cells and exhausted cells shown in average normalized intensity difference plots (Fig. 3A-B). Unstimulated cells show increased Raman intensity in peaks associated with carotenoids^26^ at 1160 cm^-1^ and 1525 cm^-1^, with heightened standard deviation at these peaks due to heterogeneity in carotenoid uptake across and within donors as shown in donor-separated Raman spectra (Supplementary Figs. 2-4). Unstimulated cells also show greater Raman intensity at nucleic acid-associated peak 798 cm^-1^ (Fig. 3A-B), reflecting more condensed chromatin due to its nondividing state.^26^ Contrastingly, activated and exhausted cells show increased Raman intensity at lipid-associated peaks at 1448 cm^-1^ and 1661 cm^-1^, as well as at protein synthesis peaks such as 1006 cm^-1^ due to greater metabolic activity.^27^ Between activated and exhausted cells, exhausted cells show greater Raman intensity at 1342 cm^-1^, 1453 cm^-1^, and 1559-1561 cm^-1^, while activated T cells have more prominent scattering at 732 cm^-1^, 787-788 cm^-1^, 817-820 cm^-1^, 1259-1263 cm^-1^, and 1275 cm^-1^. Both Raman systems reveal state-dependent differences in the phenylalanine region (1002-1011 cm^−1^), reflecting variations in protein composition and local biochemical environment. Assignment of Raman bands of interest is made in Table 1; the Raman background spectrum is shown in Supplementary Fig. 6.

**Table 1.** Raman peak assignments for cell state differentiating bands.

| Wavenumber (cm<sup>-1</sup>) | Assignment | Biological Interpretation |
| --- | --- | --- |
| 732 | Adenine | Higher nucleotide turnover in activated T cells<sup>43</sup> |
| 787-788 | Cytosine/Thymine ring breathing mode | Higher nucleotide turnover in activated T cells<sup>43</sup> |
| 798 | PO<sub>2</sub><sup>-</sup> symmetric stretching | Chromatin compacted state in unstimulated cells<sup>26,44</sup> |
| 817-820 | Phosphodiester bond in Nucleic Acids | Higher nucleotide turnover in activated T cells<sup>43</sup> |
| 1002-1011 | Phenylalanine ring breathing and C-CH<sub>3</sub> stretching | Protein synthesis, remodeling, and microenvironment shifts in activated and exhausted cells<sup>43,45-48</sup> |
| 1160 | C-C stretching vibrations of carotenoids | Lipophilic antioxidants in unstimulated T cells<sup>26,49</sup> |
| 1259-1263 | CH deformation in lipids | Unsaturated lipid remodeling in activated effector-like T cells during cell growth<sup>50</sup> |
| 1275 | Amide III | Proteome remodeling in activated T cells<sup>43,45,46</sup> |
| 1342 | CH bending/deformation in proteins | Protein aggregation in the proteotoxic stress response in exhausted T cells<sup>51</sup> |
| 1448 | CH<sub>2</sub> bending in lipids | Lipid remodeling in activated and exhausted cells<sup>27,46,50,52</sup> |
| 1453 | CH<sub>2</sub> bending in lipids | Membrane remodeling and lipid accumulation in exhausted cells<sup>46</sup> |
| 1525 | C=C stretching vibration of carotenoids | Lipophilic antioxidants in unstimulated T cells<sup>26,49</sup> |
| 1559-1561 | N-H in-plane bending and C-N stretching of Amide II | Increased protein translation activity in the proteotoxic stress response in exhausted T cells<sup>51</sup> |
| 1661 | Amide I | Protein remodeling in activated and exhausted cells<sup>43,45,46</sup> |

**Fig. 3:**
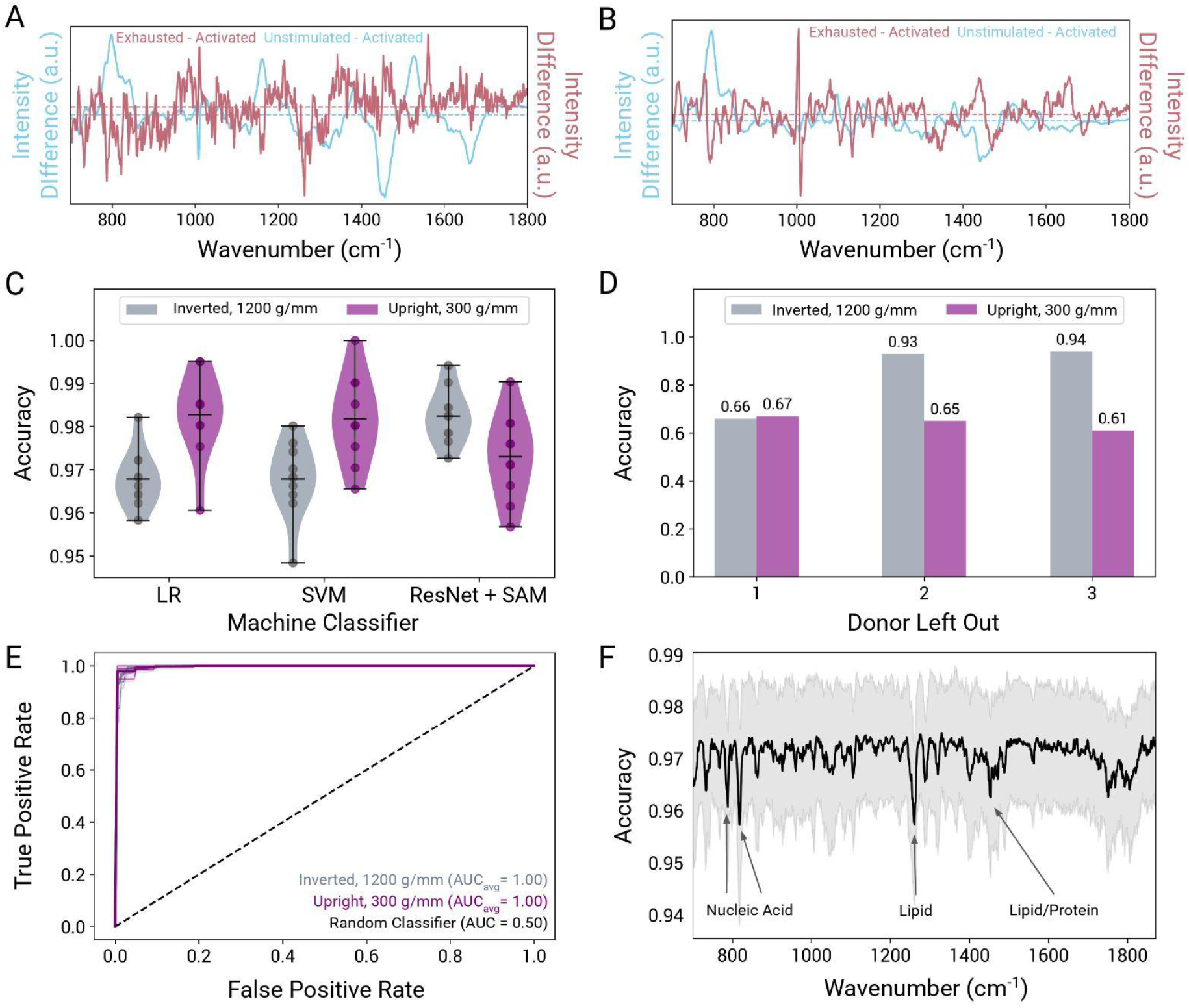
Distinguishing features and CNN classifier performance. (A-B) Raman intensity differences between cell states. Average normalized spectra in the activated cell state was subtracted from that of the exhausted or unstimulated cell state for the (A) inverted and (B) upright Raman systems. The two subtraction spectra are plotted on different y axes due to large scaling differences. Horizontal dashed lines show y = 0 for each axis. (C) Violin plots comparing accuracy for both Raman systems across three machine classifiers: LR, SVM, and our CNN, ResNet + SAM. Scatter points show each of ten splits; the mean, minimum, and maximum accuracy is indicated with horizontal lines. (D) Leave-one-donor-out cross validation (LODOCV) performed on our CNN with spectra from both Raman systems. For each donor left out of testing, the other two remaining donors were used to train the classifier. (E) Receiver operative characteristic (ROC) curve comparing classification performance between activated and exhausted cell spectra collected from the two Raman systems. Thick lines show the average across three donors with ±1 SD, and thin lines show individual donor curves. The dashed line represents a random classifier. Areas under the curve (AUC) are defined in the legend. (F) Average feature importance plot depicting the relative weights of spectral wavenumbers in the binary classification task for activated and exhausted cell spectra collected on the inverted Raman system. ±1 SD is shaded. Greater drops in accuracy indicate greater importance for classification. The four most important peaks are labeled with their vibrational mode associations.

Corresponding brightfield images (Fig. 2C and 2E) show ∼5-fold increase in cell area upon activation, contributing to success of existing size-based separation^22^ of unstimulated and activated T cells. Size and morphology of activated and exhausted T cells are more similar, necessitating a separation method with molecular-level information. Using the single cell spectra and class labels as input, we performed a stratified K-fold cross-validation of our CNN, which yielded an average classification accuracy of 98.1% and 97.4% for the inverted and upright systems, respectively (Fig. 2D and 2F). These high accuracies are maintained across donors as shown in donor-separated confusion matrices (Supplementary Figs. 2-4). As expected, the closest decision boundary occurs between the activation and exhaustion states, yet classification error remains ≤3.8% across all conditions. This small fraction of cells near the boundary is consistent with inherent heterogeneity within the cell populations at any time point. Although stimulation beads are added at a 1:1 bead-to-cell ratio, individual cells may not experience identical stimulation histories–some cells may bind to beads more than once over the four-day incubation period, while others may not bind at all. Moreover, flow cytometry-based class labels reflect population-level phenotypes rather than single cell ground truth, and no current method enables direct, non-perturbative correlation of single cell Raman spectra with fluorescent markers in suspension. Nevertheless, these results indicate the ability of Raman spectroscopy and machine learning to distinguish activated and exhausted cells with high accuracy directly from culture and without antibody labels.

We then asked whether our three-state prediction approach could be expanded to include the second stimulation at Day 8 and enable identification of cells in transition states between activation and exhaustion. Additional training and evaluation of our CNN revealed average classification accuracies of 93.4% and 92.1% for the inverted and upright systems, respectively (Supplementary Fig. 7A-B). Manufacturer instructions for Dynabeads Human T-Activator CD3/CD28 indicate that T cells may begin to exhibit signs of exhaustion starting at days 7-10 of continuous stimulation. Greater heterogeneity in T cell response at Day 8, compared to Day 4 and Day 12, likely contributes to slightly lower classification accuracies for the four-state prediction compared to the three-state prediction. Nevertheless, these results suggest that Raman spectroscopy and machine learning can be used to identify earlier indicators of T cell exhaustion, allowing for improved disease staging in chronic illness and near real-time interventions of culture conditions in cell therapy manufacturing to maintain batch quality for higher persistence and potency *in vivo*.

### CNN analysis reveals discriminating features across donors

We also examined how our CNN performed relative to other machine classifiers such as logistic regression(LR) and support vector machine (SVM). Across ten splits and both Raman instruments, all three classifiers showed average classification accuracies above 96% (Fig. 3C). The upright Raman system yielded greater variation compared to the inverted system, likely reflecting the differences in how these spectra were acquired. The inverted system collected multiple spectra at different locations within each cell which were then averaged to produce a more stable whole-cell signature. The upright system, on the other hand, collected a single spectrum per cell which is more sensitive to subcellular sampling location. This sampling approach increases within-class variability that impacts classifier performance across all models. For the inverted Raman system, our CNN showed 1.5% greater average classification accuracy across ten splits. Our CNN uses a ResNet architecture with a sharpness aware minimization (SAM) optimizer, which we have previously demonstrated to improve accuracy and generalizability for spectral classification tasks.^38^ However, for the upright Raman system, our CNN showed a slight decrease in average classification accuracy relative to LR and SVM. Because convolutional filters emphasize local peak patterns in addition to global feature weights,^39^ the CNN may be more sensitive to the subcellular heterogeneity in spectra collected on the upright system, contributing to the slight drop in performance.

This sensitivity to subcellular heterogeneity is also reflected in the LODOCV test in which we assessed the biological robustness of our CNN against new donor data. In this assessment, spectra from one of the three donors was excluded from training and used to test the resulting model, repeating this process for all donors. Fig. 3D reveals ∼20% greater classification accuracy across all donors for spectra collected on the inverted Raman system compared to the upright system, likely due to differences in spectra acquisition: whole cell signatures derived from multi-point scanning may better capture donor-specific T cell responses and improve generalizability compared to single point spectra. Average AUC for ROC curves, however, is 1 for both systems (Fig. 3E), further demonstrating our method’s sensitivity in identifying T cell exhaustion across instruments and acquisition parameters. For Donors 2 and 3, classification accuracies exceed 92% for the inverted Raman system, however, Donor 1 spectra yield ∼67% accuracy for both systems (Fig. 3D). Notably, spectra from Donors 2 and 3 were collected from cells with less than one month of cryopreservation, while Donor 1 cells had spent approximately six months in cryopreservation at the time of analysis. The effects of cryopreservation on cell phenotype have been documented extensively,^40–42^ with substantial changes to protein expression and metabolic activity observed after just one night. These shifts, indicative of cell stress, may result in Raman spectra that resemble the spectra of truly exhausted T cells. Therefore, though accuracies are high when Donor 1 spectra are included in classifier training, accuracy suffers when these spectra are excluded due to molecular-level differences in cell state compared to Donors 2 and 3.

Interestingly, the resulting confusion matrix after the LODOCV task for Donor 1 reveals that the primary source of error occurs when activated T cells are misclassified as exhausted T cells; exhausted T cells themselves are classified with high accuracy (Supplementary Fig. 8). This result could suggest that long-duration cryopreservation primarily affects T cell activation, aligning with prior studies showing impaired activation,^40^ cytokine production,^42^ and proliferation^42^ in frozen cells whose effect increases with duration. An additional possibility is that cells that have been frozen for long periods of time may be more easily exhausted with fewer stimulations. The relationship between cryopreservation and T cell exhaustion has not been extensively documented. More rigorous testing with various timepoints in the freezer should be conducted to confirm the relationship between freezing time, activation response, and exhaustability. As clinical samples and autologous cell therapies typically operate with fresh or briefly frozen cell material, we expect these cells to have similar T cell responses to Donors 2 and 3. For off-the-shelf cell therapies where cell material may be frozen for several months, however, it is recommended that additional testing with other, similarly frozen cell material take place to ensure model generalizability across donors. Additional LODOCV results for other classifiers are shown in Supplementary Fig. 9.

To identify which wavenumbers drive differentiation of activated and exhausted cells, we isolated spectra from the two states and perturbed cell spectra five times at each wavenumber using a Voigt distribution.^53^ We re-trained our CNN for the binary classification task using ten splits of the original spectra and evaluated classification accuracy with the perturbed test spectra for each of the ten splits used previously. Resulting accuracies for perturbed spectra collected on the inverted system are shown in Fig. 3F, where greater drops in accuracy indicate greater importance of the spectral feature in the classification task. The Raman peak with the greatest drop in accuracy is 1260 cm^-1^, followed by 788 cm^-1^, 819 cm^-1^, and 1457 cm^-1^. These lipid- and nucleic acid-associated peaks align with the previously identified peaks in Table 1 that are elevated in either activated or exhausted cells, confirming the model’s biological relevance in learning molecular differences that separate the two cell states. These Raman peaks are similarly identified in the feature importance plots of the other machine classifiers (Supplementary Fig. 9), along with 863 cm^-1^ and 1659 cm^-1^. Overall, lower drops in accuracy for each feature are observed when perturbed spectra are input to our CNN compared to LR and SVM, demonstrating increased robustness against small changes in peak intensity likely due to the SAM optimizer which encourages learning broader, more stable spectral patterns.^38^ Similar features are identified in the feature importance plots for spectra collected on the upright system shown in Supplementary Fig. 10, along with two new, highly important features: 1002 cm^-1^ and 1011 cm^-1^. This result matches observations in the intensity difference plots (Fig. 3B) and reflects changes in phenylalanine due to altered protein synthesis and remodeling. Differences in feature extraction between the two Raman systems are likely due to the aforementioned differences in spectral collection, where the inverted system provides a global biochemical readout and the upright system samples a single intracellular location. Therefore, features identified with the upright system may reflect localized contributions from subcellular regions associated with exhaustion that may not hold as much weight in the whole-cell readout from the inverted system.

### Exhaustion percentage is predicted in mixed cell batches

Because real clinical samples often contain mixtures of cell states at unknown proportions, we assessed whether our model could accurately infer the percentage of each cell state within heterogeneous populations. To do this, we systematically designed an experiment of 100 cell batches with a randomly selected percentage (X%) of exhausted cell spectra. The remaining percentage was split randomly between activated (Y%) and unstimulated (Z%) cell spectra. These spectra were then randomly selected from the total number of available cell spectra for each class for each donor. These donor-specific batches, each with 100 cell spectra, were then used to test our CNN that had been pre-trained with spectra from the other donors (Fig. 4A). With spectra acquired using the inverted Raman system, predicted percentages of exhausted, activated, and unstimulated cells showed strong linear relationships to the ground truth percentages of each class with R^2^ values of 1, 0.94 and 1, respectively (Fig. 4B). Small deviation in the prediction of activated or exhausted cell states could be caused by inherent heterogeneity in the cell populations due to non-identifical stimulation histories or by donor-specific responses to activation and exhaustion. The same analysis for spectra collected on the upright Raman system yielded poor R^2^ values (Fig. 4D), further emphasizing the importance of an averaged whole-cell readout rather than a single point spectrum.

**Fig. 4:**
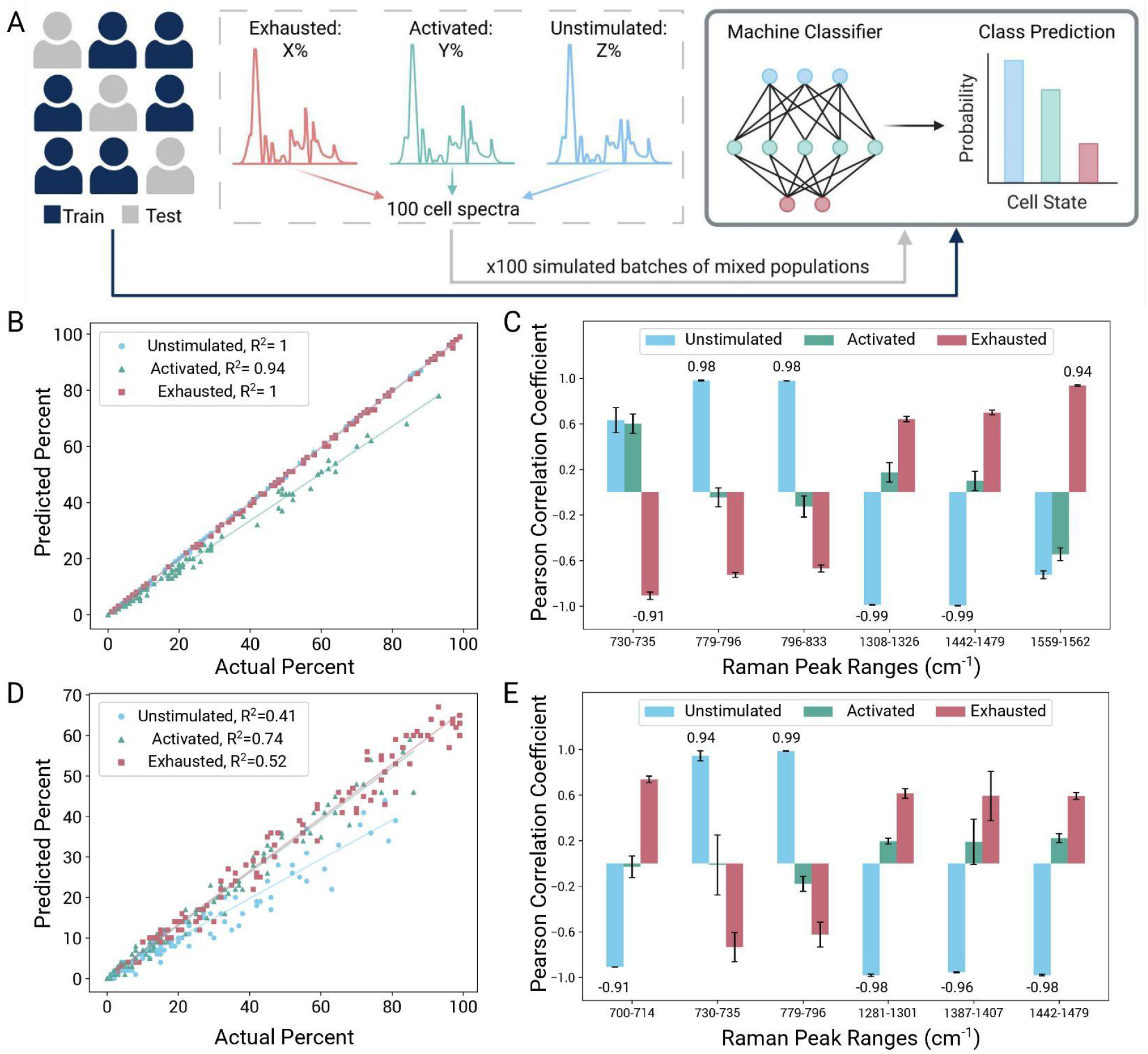
Mixed population testing. (A) Schematic of mixed population generation from single cell spectra. For each donor, 100 cell batches of mixed populations were generated by pooling 100 randomly sampled spectra according to randomly selected percentages from each cell state. These spectra batches were then used to test our CNN which had been previously trained on cell spectra from the other donors. The CNN assigns class labels to each cell spectra and the predicted percentage of each cell state is compared to the ground truth compositions of each batch. (B) Representative scatter plot for Donor 3 showing the relationship between the predicted and actual percentage of each class with spectra acquired from the inverted Raman system. Each scatter point represents a single batch used to test the CNN. The best fit lines are plotted for each cell state. Prediction R^2^ values defined in the legend indicate how well our CNN predicted the actual percentages. (C) Grouped bar plot showing the Pearson correlation coefficients between the ground truth percent of each cell state and the average AUC for select Raman peak ranges in normalized spectra batches. Correlation coefficients were averaged for Donors 2 and 3, and ±1 SD is indicated with error bars. (D-E) Corresponding analyses to (B–C), conducted with spectra acquired from the upright Raman system.

Unique to exhausted T cells, average normalized Raman spectra for each batch show decreasing AUC at adenine peak region 730-735 cm^-1^ and increasing AUC at amide II protein peak region 1559-1562 cm^-1^ (Fig. 4C) with Pearson correlation coefficients of −0.91 and 0.94, respectively. Interestingly, few peak ranges were found to be highly correlated with T cell activation; most were correlated with either the unstimulated or exhausted state (Fig. 4C and 4E). One explanation is that T cell activation is a more dynamic state with T cell response having greater heterogeneity even within donors. Unstimulated T cells may be more uniform in their molecular composition, and although T cell exhaustion is progressive, our T cells may have arrived at a more uniform state of terminal exhaustion by Day 12. This heterogeneity in T cell activation supports the use of advanced machine classifiers like our CNN for label-free cell state identification, rather than regression-based models trained with Raman peak AUCs alone. Our CNN learns features from the whole spectra, including individual peak intensities and widths, peak ratios, peak shoulders, etc., enabling improved generalization to heterogeneous cell states and samples. While our mixed batches were generated by pooling known percentages of spectra from each cell state, this approach allowed us to determine our method’s robustness against heterogeneous mixtures without introducing confounding effects such as pipetting or sampling errors which could result in incorrect percentage labels. Altogether, these findings demonstrate that our model can reliably navigate the intrinsic variability of T cell samples, providing meaningful feature interpretation for deployment in clinical and laboratory settings.

## Outlook

In summary, we demonstrate a robust and accurate methodology to distinguish exhausted T cells from activated and unstimulated T cells directly from culture and without antibody labels, destructive processes, or days-long functional assays. Unlike the flow cytometry gold standard, our Raman-based assessment is not limited to a set of protein markers and does not require time-consuming and tedious sample preparation steps like washing, labeling, or fixation. Raman generates a complete molecular profile of T cell exhaustion with discriminative features often excluded from flow cytometry analysis. Our approach is robust across different Raman instruments and acquisition parameters, with multi-point whole-cell spectra providing the highest accuracy when evaluating donors not included in training. Our custom CNN classifier ResNet + SAM outperforms commonly used models and learns biologically relevant spectral features, distinguishing T cell activation and exhaustion primarily with Raman peak regions associated with nucleotide turnover, lipid remodeling, and amide II protein alterations. Moreover, the accuracy of our methodology is conserved in heterogeneous cell batches, where the percentage of exhausted T cells strongly correlates to the AUC of distinguishing peak regions. These results support our method’s usability in applications such as immune diagnostics, cell therapy manufacturing, and therapy monitoring where T cell state heterogeneity is expected. Notably, our method also detects transition states where activated T cells begin to present signs of exhaustion, enabling near real-time interventions to prevent further exhaustion and restore immune function.

Further work is needed to characterize the molecular fingerprints of cells that have been cryopreserved for extended periods, an increasingly relevant scenario as allogeneic cell therapies expand. Additionally, the development of methods to allow one-to-one pairing of cell phenotype information across modalities without perturbative immobilization would help capture inherent heterogeneity and further strengthen model performance. In its current form, our method provides valuable information on T cell quality and can be integrated into existing workflows for at-line or off-line monitoring of T cell exhaustion during culture and clinical assessment of T cell state for disease diagnosis and treatment selection. With advances in Raman-activated cell sorting,^31–33^ this work could also support purification of starting cell material in cell therapy manufacturing and expand treatment options for millions of patients suffering from chronic diseases like HIV, tuberculosis, malaria, and cancer.

## Methods

### T cell isolation and stimulation

Pan T cells from three healthy donors were isolated from frozen peripheral blood mononuclear cells collected from fresh Leukopaks (STEMCELL Technologies) using the EasySep Human T Cell Isolation Kit (STEMCELL Technologies, Cat. # 17951). The T cells were then allowed to rest for one day in 6-well plate cultures (Thermo Fisher Scientific, Cat. # FB012927) in Roswell Park Memorial Institute (RPMI) 1640 medium (Thermo Fisher Scientific, Cat. # 11-875-093) complete with 10% Fetal Bovine Serum (FBS, VWR, Cat. # 89510-188) and 1% penicillin-streptomycin (Thermo Fisher Scientific, Cat. # 15140148) with 30 U/mL human rIL-2 (Thermo Fisher Scientific, Cat. # PHC0026). The T cells were then activated using Dynabeads Human T-Activator CD3/CD28 (Thermo Fisher Scientific, Cat. # 11131D) at a 1:1 bead-to-cell ratio. For maximum cell recovery, cultures were maintained for four days before bead removal with the EasySep Magnet (StemCell Technologies, Cat. # 18000) and cell harvest for analysis or sequential stimulation. Cells were re-stimulated twice with fresh beads, media, and rIL-2 for a total stimulation time of twelve days in an incubator at 37°C with 5% CO_2_.

### Flow cytometry

To obtain cell state labels at the population level, flow cytometry was performed with a BD FACSymphony™ A3 Cell Analyzer with the BD FACSDiva software. Pre-diluted mouse anti-human antibodies and catalog numbers are detailed in Supplementary Table 1 and used according to manufacturer instructions. Following magnetic bead removal, cells were washed with wash buffer, phosphate-buffered saline (PBS) with 0.1% sodium azide, and resuspended in staining buffer (PBS with 0.1% sodium azide + 2% FBS) prior to incubation with antibody cocktail for 30 min at 4°C in the dark. Cells were washed once more with wash buffer, resuspended in staining buffer, and strained using a 35 μm filter capped tube (STEMCELL Technologies, Cat. # 100-0087) prior to analysis. Compensation was performed using the BD™ CompBeads Set Anti-Mouse Ig, κ (BD Biosciences, Cat. # 552843). Analysis was conducted in FlowJo v10 using the gating strategy shown in Supplementary Fig. 1.

### Raman spectroscopy

Following magnetic bead removal, single cell suspensions were plated with fresh media without rIL-2 into quartz-bottom petri dishes (Waken B. Tech, Cat. # SF-S-D12). To demonstrate robustness against instrument-to-instrument variability and spectral resolution, single cell spectra were collected either with a commercial, upright (WITec Alpha300) or custom-built^25^, inverted Raman microscope, both equipped with a 785 nm laser. For the commercial, upright microscope, a silicon wafer was set underneath the quartz-bottom petri dish for better visualization of cells; the top of the dish was removed to allow sufficient access for focusing.

Single cells were illuminated for 15 s with 2 accumulations at 50 mW power using a 50× dry objective. A single point spectra was collected for each cell with a 300/mm grating. For the custom-built, inverted microscope,^25^ quartz-bottom petri dishes were placed onto a sample holder above a 60× water immersion objective and encased in an on-stage incubator set to 37°C with 5% CO_2_. Multiple point spectra per cell were recorded using a graphical user interface between *napari* and *Micro-manager* powered by *pymmcore-plus*. Point spectra were collected with an exposure time of 350 ms and 258 mW power at the sample with a 1200 g/mm grating and averaged to obtain single cell spectra after cosmic ray and fluorescence removal. The total number of single cell spectra for unstimulated, activated and exhausted cells was 1930, 1535, and 1570 for the inverted system and 567, 713, and 749 for the upright system. The total number of cell spectra for the second stimulation at Day 8 was 1572 and 691 for the inverted and upright systems, respectively. Total number of single cell spectra for Donors 1, 2, and 3 was 512, 756, and 761 for the upright system and 2636, 1330, and 1069 for the inverted system.

### Spectra pre-processing

The spectral fingerprint region (700 cm^-1^ to 1800 cm^-1^) was subjected to cosmic ray removal,^54^ where peaks with a z-score threshold above 5 are replaced with the mean intensity of non-peak values within a window size of 10. Autofluorescence removal was performed using alternating least squares in the baseline function in *rampy*.^55^ The spectra was then smoothed using a Savitzky-Golay filter with a third-order polynomial and a window length of five data points using the *savgol_filter* function from *scipy*. For spectra visualization and intensity comparison, the smoothed spectra were subjected to Standard Normal Variate (SNV) normalization. For input to machine classifiers, the smoothed spectra was instead standardized to a mean of 0 and a standard deviation of 1 using the *StandardScaler* function from *scikit-learn*.

### Spectra classification and feature importance

To classify single cell spectra according to stimulation number and cell state, single cell spectra and class labels were used to train and evaluate three machine classifiers: LR and SVM from *scikit-learn* and our CNN adapted from a ResNet architecture.^56^ The SAM optimizer^38^ was used instead of Adam with a Scaled Exponential Linear Unit (SELU) activation function and a batch size of 16. To assess the classification performance, we used stratified K-fold cross-validation across ten splits (*scikit-learn StratifiedKFold*). For each split, the train set is used to fit the *StandardScaler*, and the scaler transforms both the train set and the test set. The model is then trained on the scaled train set and evaluated on the scaled test set. In this way, we generate unbiased predictions for all data, and these are compared against the class labels to generate confusion matrices and the overall accuracy (*skikit-learn, accuracy_score* and *confusion_matrix)*. For the LODOCV task, spectra from one donor were selected as the test set, while the remaining two donors were used as the train set. Model training and evaluation was repeated for all three donors to assess model rigor against biological variability.

To determine the most important spectral features driving classification performance between the activated and exhausted T cell spectra, we repeated K-fold cross-validation across ten splits for the binary classification task. For each split, we perturbed test set spectra prior to evaluating classification using models trained on the unperturbed train set. We generated five perturbations for each probing wavenumber by using a Voigt distribution^53^ created by random sampling of spectral intensity values at that wavenumber. We evaluate classification accuracy for each perturbation and split and average the resulting accuracies per wavenumber; greater decreases in accuracy after perturbation signify greater importance of that spectral feature. Using the binary spectra, ROC curves were generated from classifier-defined probabilities for the exhausted state across ten train-test splits.

### Mixed batch testing

Heterogeneous T cell batches with varying exhaustion composition were generated by randomly selecting a percentage for exhausted cell spectra. The remaining percentage was split randomly between unstimulated and activated cell spectra. These spectra were then randomly selected from the total number of available cell spectra for each class for each donor at the designated percentages. These donor-specific batches, each with 100 distinct cell spectra, were then used to test our CNN that had been pre-trained with spectra from the other donors. This process was repeated for a total of 100 batches per donor. R^2^ values were determined using the *r2_score* function from *scikit-learn*. Pearson correlation coefficients (two-sided) were calculated using the *pearsonr* function in *scipy* to determine the relationship between the average AUC of Raman peak regions of interest in SNV normalized spectra and the percentage of each cell state in the cell batch.

## Data Availability

The flow cytometry data, raw and processed spectra, and analysis scripts may be accessed upon request from the corresponding authors.

## Acknowledgements

This work was supported in part by the National Science Foundation Graduate Research Fellowship and the Advanced Undergraduate Research Opportunities Program (SuperUROP) at MIT. We would like to thank Dr. Koseki Kobayashi-Kirschvink and Dr. Jeon Woong Kang for their support in part of this work conducted in the MIT Laser Biomedical Research Center. We also thank Dr. Ian Hunt-Isaak from the Hekstra lab at Harvard University for his support in streamlining multi-point spectra acquisition in suspension cells. Finally, we thank the Koch Institute’s Robert A. Swanson (1969) Biotechnology Center for technical support, specifically the flow cytometry core facility.

## Declaration of Competing Interest

MM, SP, and LFT are co-inventors on a patent application related to the label-free identification of cell state characteristics. SR declares no conflict of interest.

## Author Contributions

**MM**: writing (original draft), writing (review and editing), conceptualization, investigation, visualization, formal analysis; **SP**: writing (original draft), writing (review and editing), investigation, formal analysis; **SR**: investigation; **LFT**: writing (review and editing), resources, supervision, project administration, conceptualization

## Notes

### Competing Interest Statement

The authors have declared no competing interest.

