## Supplemental Materials for "Nucleic acid turnover and lipid remodeling distinguish T cell activation and exhaustion states via label-free Raman spectroscopy"

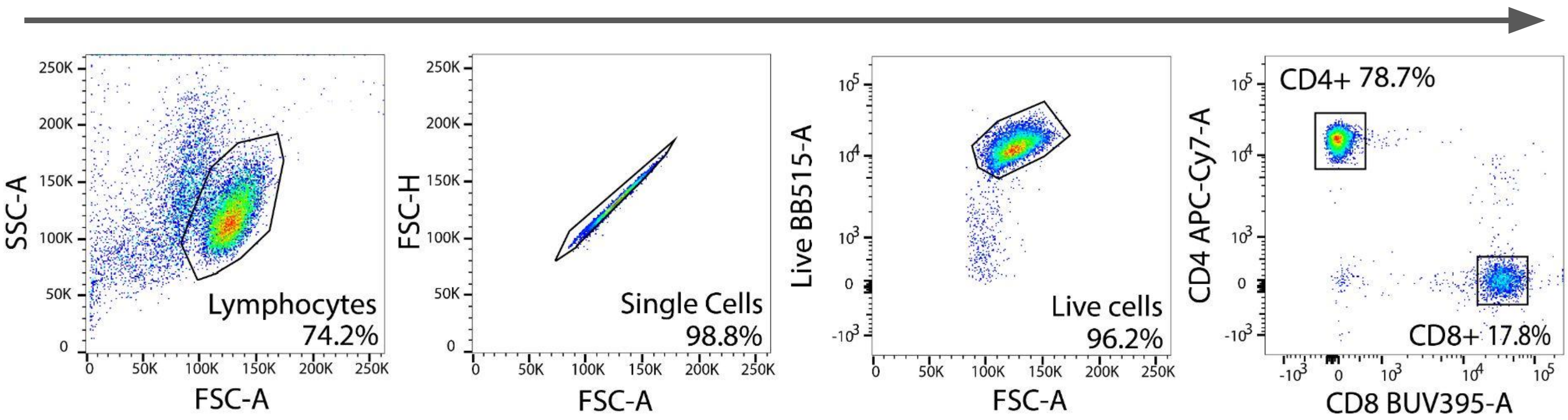


**Supplementary Fig. 1.** Gating strategy for flow cytometry analysis

**Supplementary Table 1.** Flow cytometry markers

| Marker | Stain | Cat. # (Supplier) |
|---|---|---|
| Live | FITC | R37601 (Fisher Scientific) |
| CD4 | APC-Cy7 | 566319 (BD Biosciences) |
| CD8 | BUV395 | 563795 (BD Biosciences) |
| PD-1 | Alexa Fluor 647 (APC) | 566850 (BD Biosciences) |
| LAG-3 | PE | 565616 (BD Biosciences) |
| TIM-3 | BV421 | 565562 (BD Biosciences) |

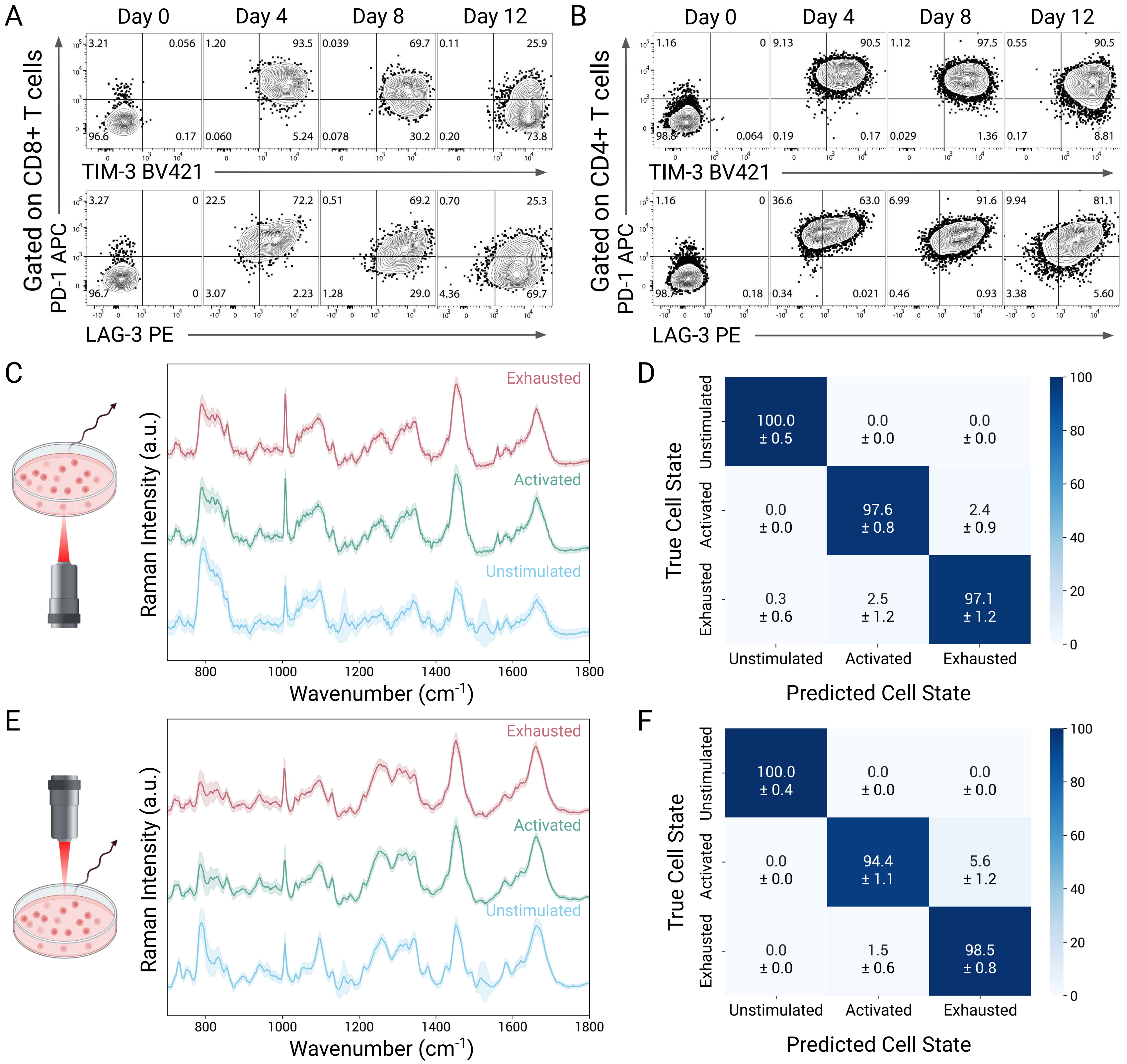


**Supplementary Fig. 2.** Raman-based identification of T cell exhaustion for Donor 1. (A-B) Representative flow cytometry analysis of (A) CD8+ and (B) CD4+ T cells for the expression of inhibitory receptors PD-1, LAG-3, and TIM-3. (C-D) Identification via the inverted Raman system (1200 g/mm grating, multi-point scanning): (C) Average normalized single cell spectra from live T cells. Spectra are grouped by cell state with ±1 standard deviations (SD) shaded and used to generate the (D) normalized confusion matrix from our CNN. (E-F) Identification via the upright Raman system (300 g/mm grating, single-point acquisition): (E) Average normalized single cell spectra from live T cells. Spectra are grouped by cell state with ±1 SD shaded and used to generate the (F) normalized confusion matrix from our CNN.

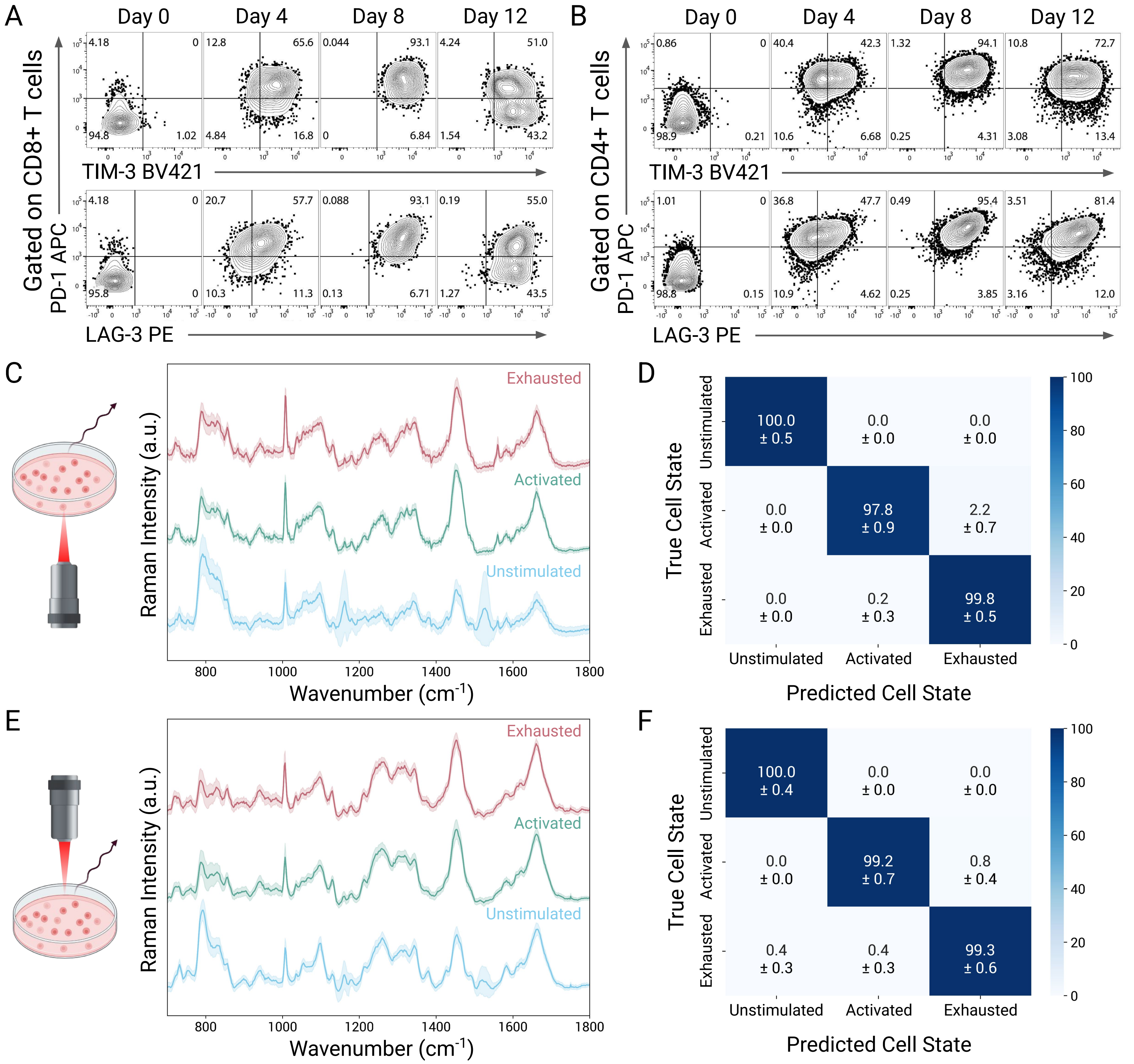


**Supplementary Fig. 3.** Raman-based identification of T cell exhaustion for Donor 2. (A-B) Representative flow cytometry analysis of (A) CD8+ and (B) CD4+ T cells for the expression of inhibitory receptors PD-1, LAG-3, and TIM-3. (C-D) Identification via the inverted Raman system (1200 g/mm grating, multi-point scanning): (C) Average normalized single cell spectra from live T cells. Spectra are grouped by cell state with ±1 SD shaded and used to generate the (D) normalized confusion matrix from our CNN. (E-F) Identification via the upright Raman system (300 g/mm grating, single-point acquisition): (E) Average normalized single cell spectra from live T cells. Spectra are grouped by cell state with ±1 SD shaded and used to generate the (F) normalized confusion matrix from our CNN.

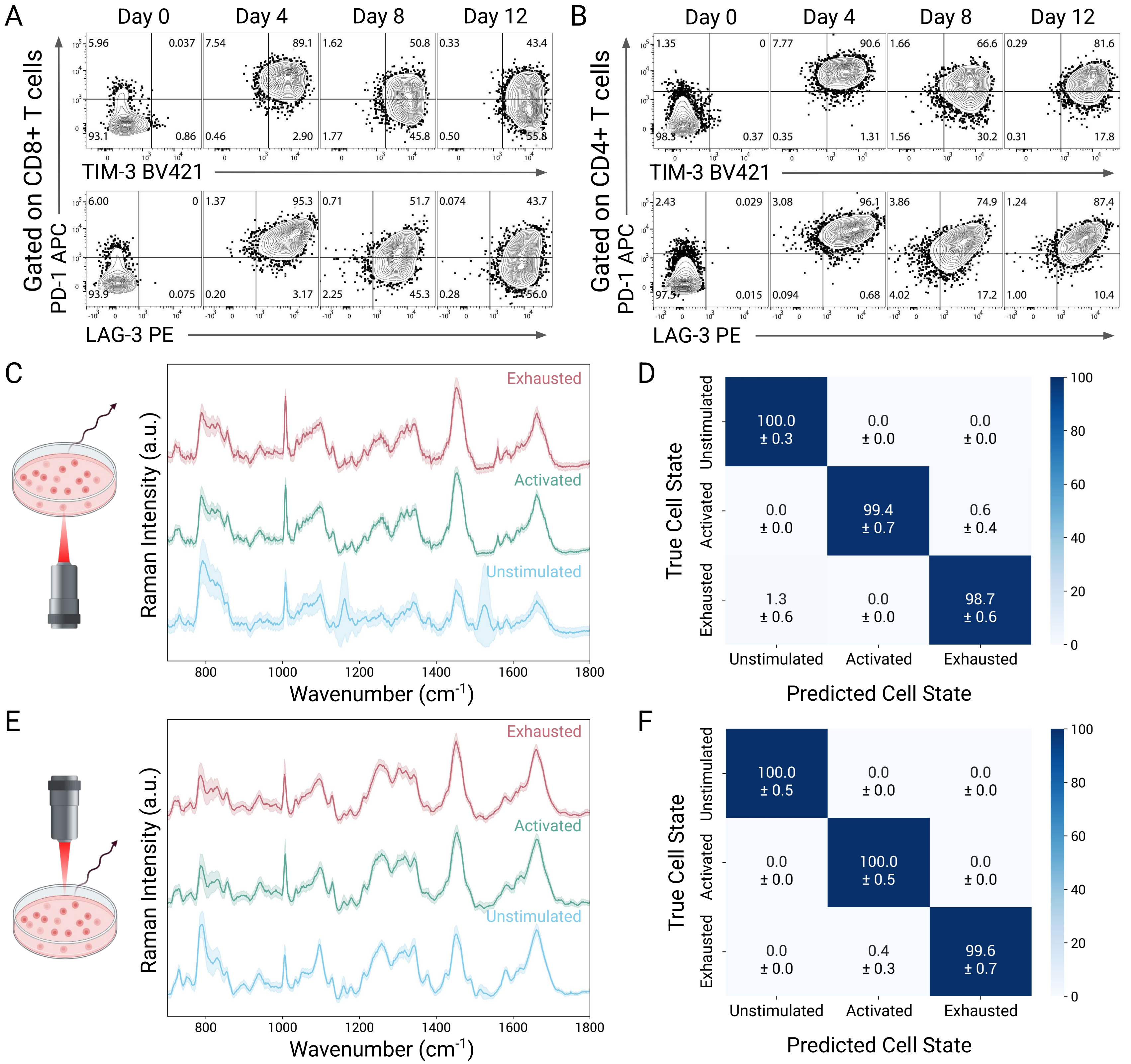


**Supplementary Fig. 4.** Raman-based identification of T cell exhaustion for Donor 3. (A-B) Representative flow cytometry analysis of (A) CD8+ and (B) CD4+ T cells for the expression of inhibitory receptors PD-1, LAG-3, and TIM-3. (C-D) Identification via the inverted Raman system (1200 g/mm grating, multi-point scanning): (C) Average normalized single cell spectra from live T cells. Spectra are grouped by cell state with ±1 SD shaded and used to generate the (D) normalized confusion matrix from our CNN. (E-F) Identification via the upright Raman system (300 g/mm grating, single-point acquisition): (E) Average normalized single cell spectra from live T cells. Spectra are grouped by cell state with ±1 SD shaded and used to generate the (F) normalized confusion matrix from our CNN.

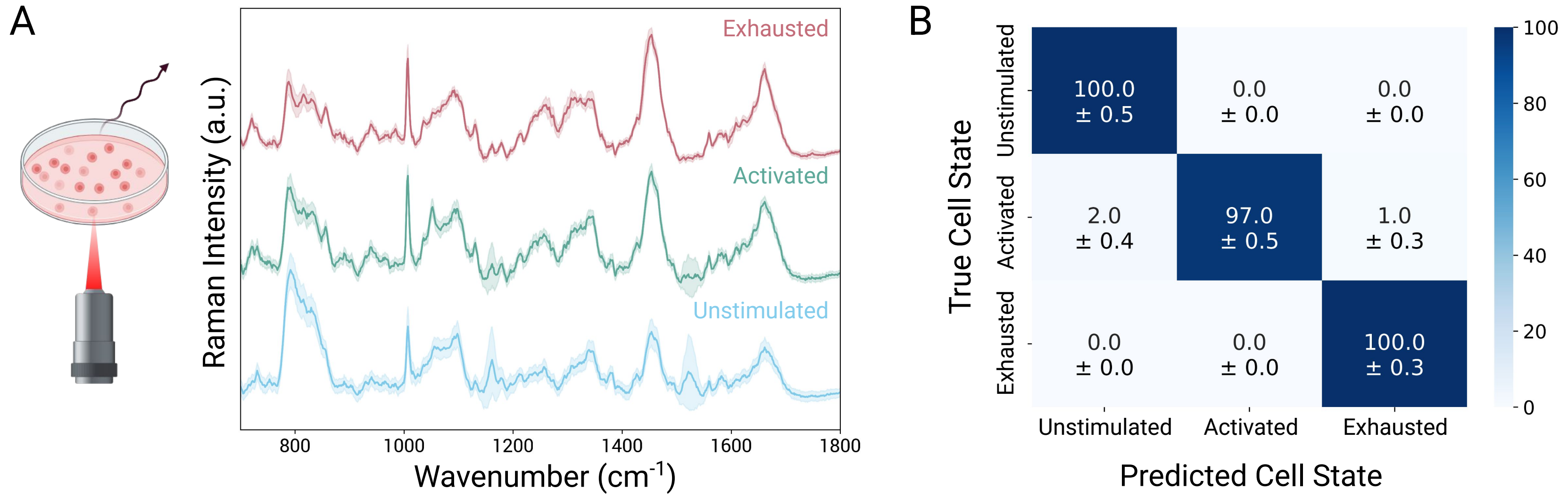


**Supplementary Fig. 5.** Raman-based identification of T cell exhaustion in CD8+ T cells alone. (A) Average normalized single cell spectra from live T cells (n=1006). Spectra were collected using the inverted Raman system (1200 g/mm grating, multi-point scanning) and are grouped by cell state with ±1 SD shaded. (B) Normalized confusion matrix generated using our CNN classifier. Performance was evaluated with a stratified K-fold cross-validation across ten splits.

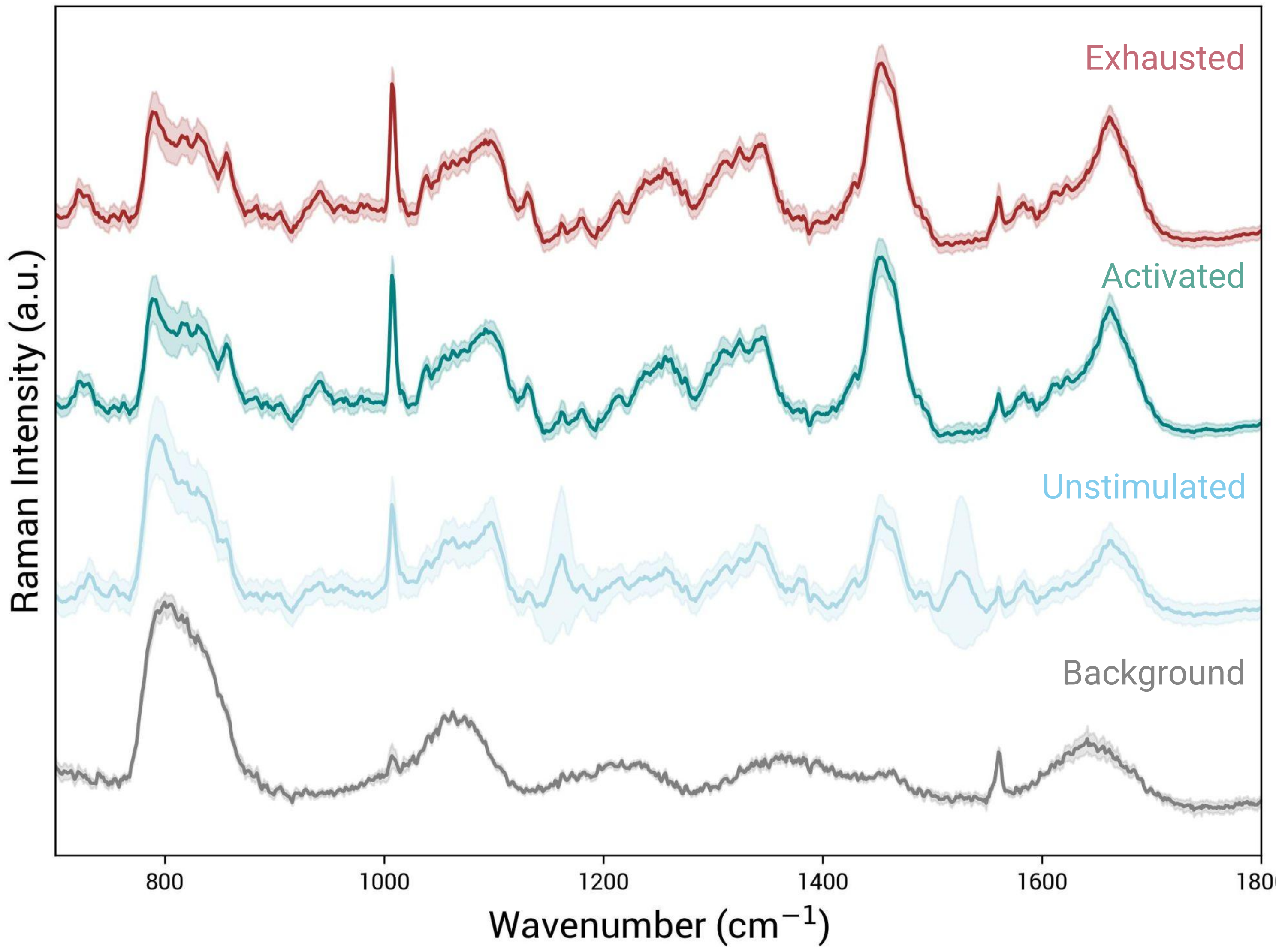


**Supplementary Fig. 6.** Average normalized Raman spectra of the background, including media and quartz bottom petri dish, in comparison to the cell spectra. Spectra were collected using the inverted Raman system (1200 g/mm grating, multi-point scanning), and ±1 SD is shaded.

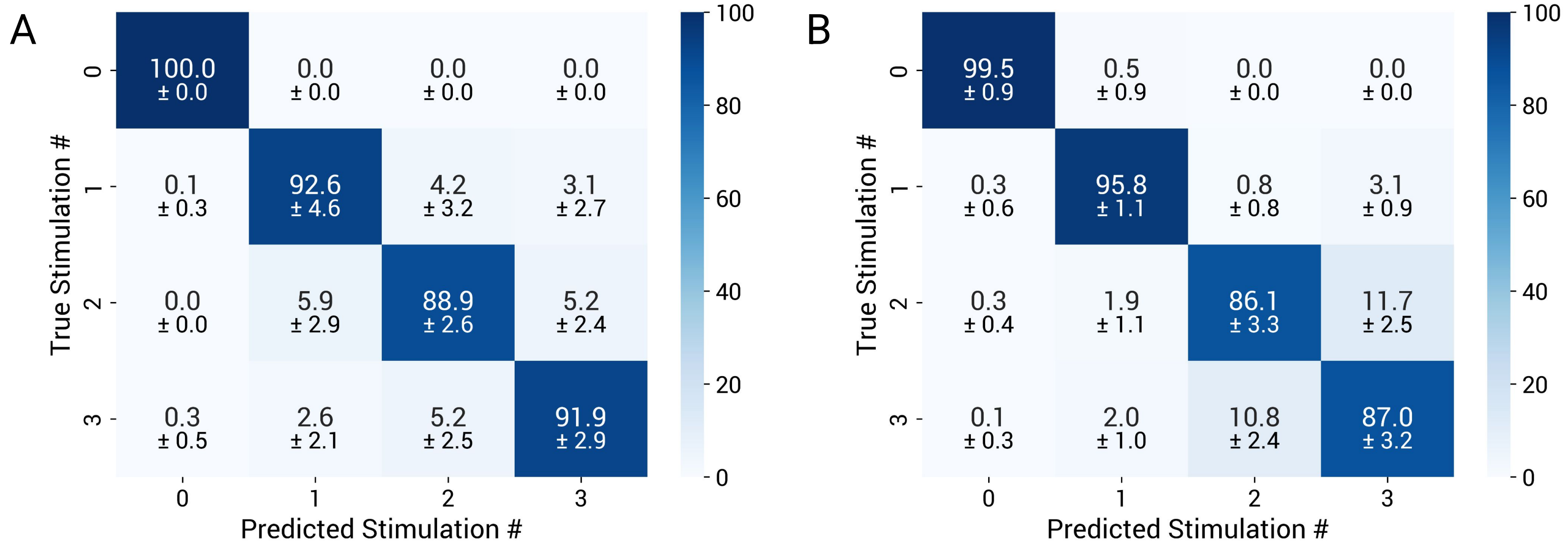


**Supplementary Fig. 7.** Four-class classification of cell spectra measured after each stimulation. Normalized confusion matrix generated using our CNN classifier on the single cell spectra collected via the (A) inverted or (B) upright Raman system. Performance was evaluated with a stratified K-fold cross-validation using ten splits across three donors with three technical replicates each.

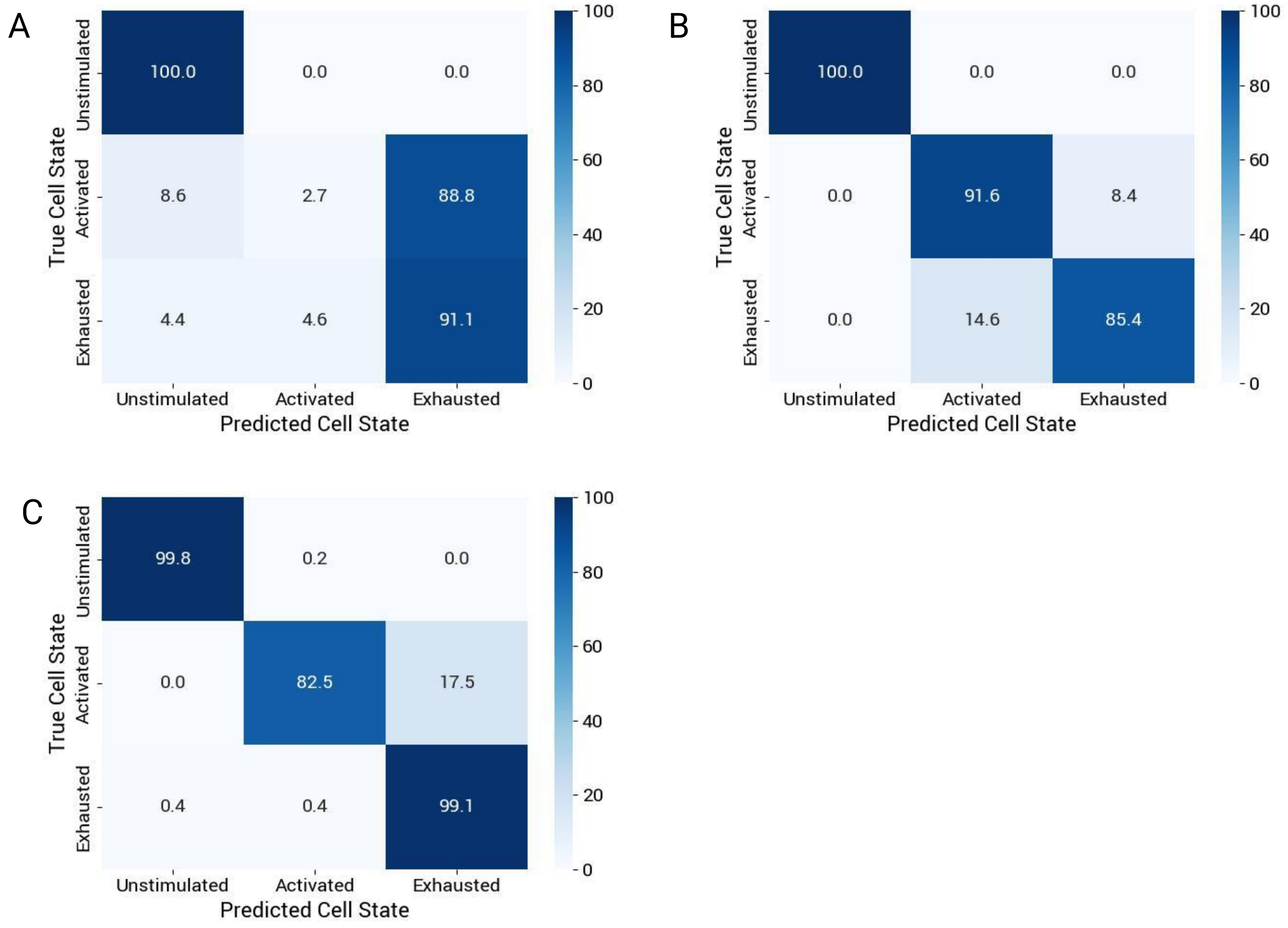


**Supplementary Fig. 8.** Normalized confusion matrices generated from cell spectra collected on the inverted Raman system from (A) Donor 1, (B) Donor 2, and (C) Donor 3 during the leave-one-donor-out cross validation task.

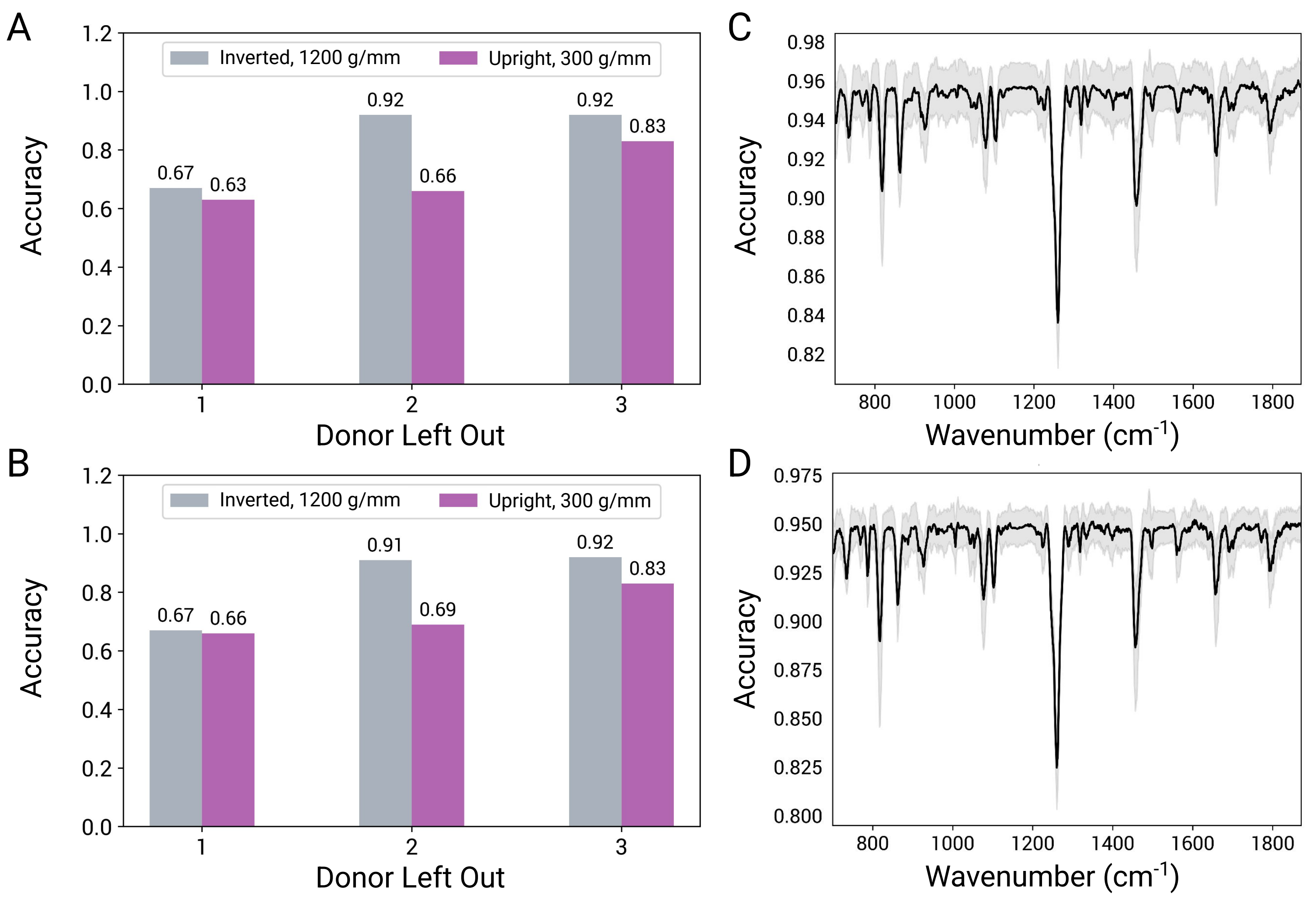


**Supplementary Fig. 9.** Machine classifier performance of logistic regression (LR) and support vector machine (SVM). (A-B) Leave-one-donor-out cross-validation performed on (A) LR and (B) SVM with spectra from both Raman systems. For each donor left out for testing, the other two remaining donors were used to train the classifier. (C-D) Feature extraction performed on (C) LR and (D) SVM to determine the relative weight of spectral wavenumber in the classification task on spectra collected via the inverted Raman system. Greater drops in accuracy indicate greater importance for classification. ±1 SD is shaded.

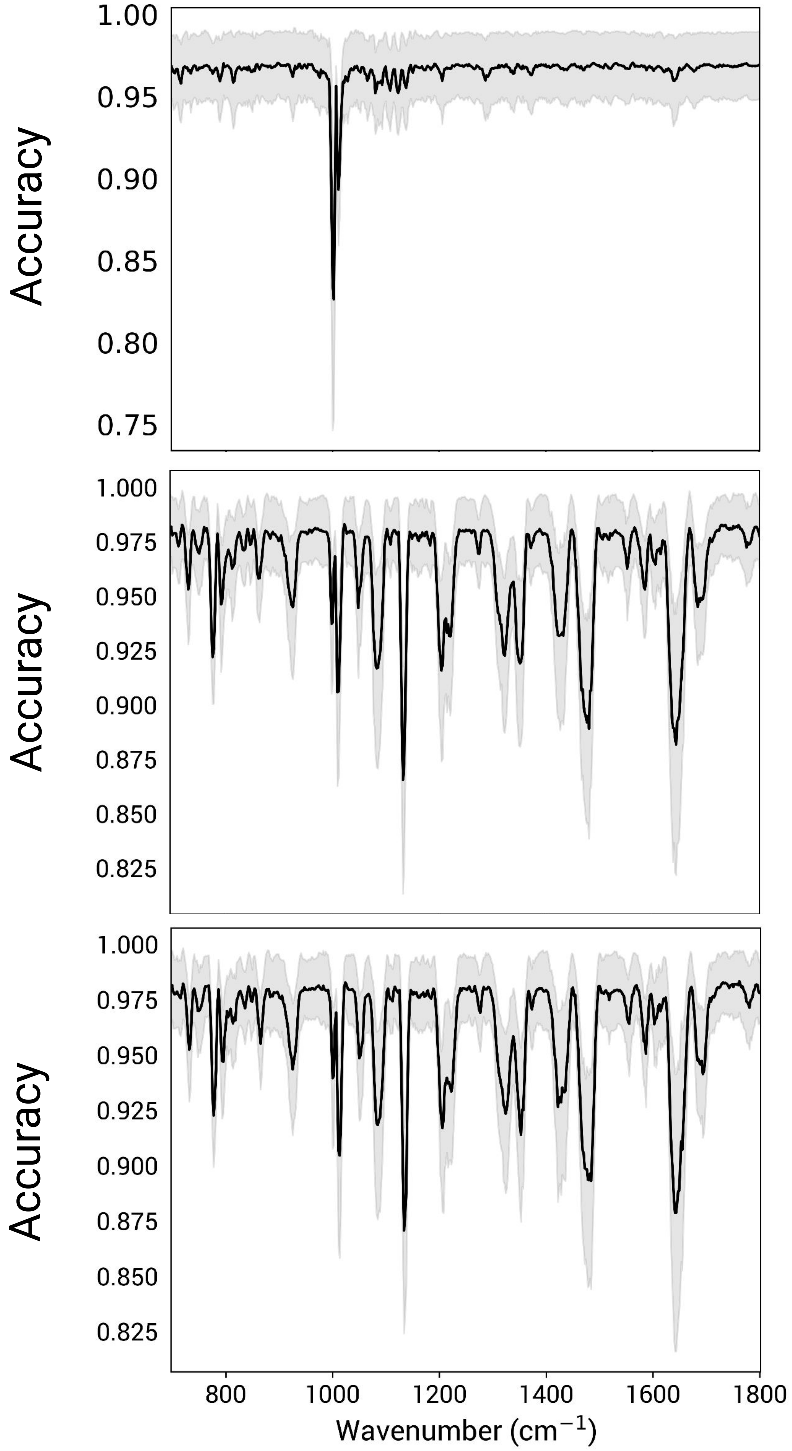


**Supplementary Fig. 10.** Feature extraction performed on (top) our CNN, (middle) SVM, and (bottom) LR to determine the relative weight of spectral wavenumber in the classification task on spectra collected via the upright Raman system. Greater drops in accuracy indicate greater importance for classification. ±1 SD is shaded.

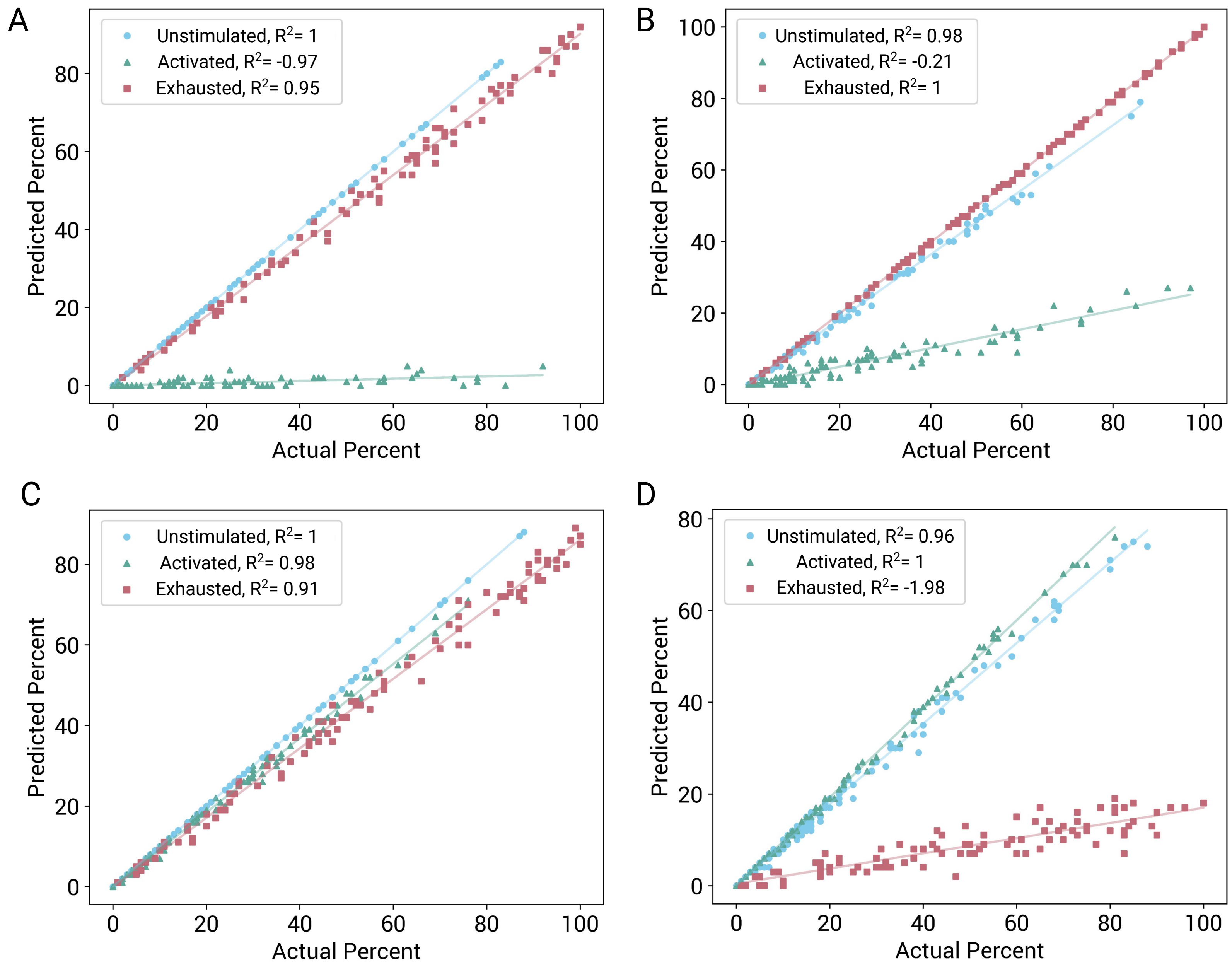


**Supplementary Fig. 11.** Mixed population testing for Donor 1 (A-B) with spectra collected on the inverted (A) and upright (B) system and Donor 2 (C-D) with spectra collected on the inverted (C) and upright (D) system. Scatter plots show the linear relationship between the predicted percentage of each class and the actual percentage of each class determined when generating the cell batches. Each scatter point represents a single batch used to test the CNN. The best fit lines are plotted for each cell state. Prediction $R^2$ values defined in the legend indicate how well our CNN predicted the actual percentages.
